# Expansion Revealing of Pathology Resolves Nanostructures Associated with Inflammatory Phenotypes in COVID-19 Decedent Human Brain Tissue

**DOI:** 10.64898/2026.05.14.725177

**Authors:** Alice E. Stanton, Jinyoung Kang, Joel W. Blanchard, Carles A. Boix, Margaret E. Schroeder, Youngmi Lee, Hanquan Su, Shiwei Wang, Eunah Yu, Amauche Emenari, Zhuyu Peng, Emre Agbas, Oyku Cerit, Demian Park, Ruihan Zhang, David A. Bennett, Peng Yin, Manolis Kellis, Robert Langer, Edward Boyden, Li-Huei Tsai

## Abstract

Expansion revealing (ExR) elucidates cellular organization by separating proteins within dense nanostructures by 20x linear expansion, but requires fixation procedures incompatible with human pathology specimens. Here, we report ExR of pathology (ExRPath), which attains ∼20 nm resolution and decrowding of such tissues, through iterative 20x expansion, adapted to human brain pathology specimens. We also report a single-shot 15x expansion protocol for such tissues (15ExMPath), achieved through one-shot 15x expansion. Applying ExRPath and 15ExMPath to COVID-19-decedent brain tissue reveals periodic amyloid nanoclusters that co-localize with SARS-CoV-2 in a rare minority of patient specimens, pointing to a potential neuroinflammatory phenotype associated with COVID-19, and highlighting the power of high-throughput nanoimaging, empowered by expansion microscopy, for discovering potential novel disease mechanisms.

---

COVID-19 is associated with acute and chronic neurological complications including cognitive fog, ischemia, headache, and dizziness {Ellul 2020}. SARS-CoV-2 virus has been detected in the brain {Stein 2022}. However, our understanding of COVID-19 neurological mechanisms and effects remains limited given that the interactions between brain proteins and viral proteins involve nanoscale biomolecules, and occur at hard-to-observe nanoscale distances. Ideally, we could visualize biomolecules associated with SARS-CoV-2, and the biomolecules of the brain, with nanoscale precision. This would allow us to hypothesize and test which brain proteins interact with viral proteins, with potential causal influence on pathological outcomes.

Conventional microscopy is unable to resolve protein-protein relationships below 200 nm, making it difficult to visualize putative interactions of viral proteins with brain proteins and other biomolecules. We recently developed expansion revealing (ExR) {Sarkar 2022}, a novel expansion-based technology that enables super-resolution imaging of brain tissue on conventional microscopes through iterative 20x physical expansion of the tissue. The resulting expansion revealed previously hidden brain nanostructures by decrowding biomolecules from each other, which in turn enabled better post-expansion staining – in some cases, making molecules visible by facilitating their labeling with fluorescent antibodies that were too large to reach the molecules when the brain tissue was in the non-expanded state. However, ExR previously could not be performed on human brain tissue, due to its requirement for fixative cocktails rarely used in standard human pathology, limiting the use of this technology in analyzing post-mortem human brain samples or other human pathology specimens.

To overcome this limitation, we invented a new approach called expansion revealing of pathology (ExRPath). ExRPath expands conventionally fixed human (and mouse) brain tissue with the classical ExR polymer, which achieves 20x linear expansion through iterative application of 4.5x expansion gels. This protocol separates proteins from each other, facilitating immunolabeling and imaging on a conventional microscope with ∼20 nm precision. We also recently developed a single-shot 20x expansion (20ExM) protocol {Wang 2024}, which achieves super-resolution imaging of brain tissue on conventional microscopes through one round of 20x physical expansion. Both ExR and 20ExM have decrowding and 20x physical expansion capabilities; therefore, we further optimized the 20ExM protocol for human pathology brain specimens, ultimately achieving 15x expansion by a one round procedure (which we call 15ExMPath).

ExRPath differs from ExR in the fixative, protein anchoring, and denaturation steps. ExR starts with cardiac perfusion of mice or the treatment of cultured neurons using paraformaldehyde (PFA) and acrylamide, whereas ExRPath starts with formalin-fixed human tissues or PFA cardiac perfusion in mice. ExRPath requires the specimen to be treated with Acryloyl-X SE (AcX) to anchor proteins to the hydrogel, and protein softening at 121°C and 30.5 psi in denaturation buffer for sufficient softening with long-term fixed human brain specimens. The 15ExMPath procedure differs from 20ExM in its protein anchoring step, using AcX treatment as in ExRPath instead of N-acryloxysuccinimide (AX), and in its protein softening step, which consists of overnight incubation at 70 °C followed by 60 min at 121 °C and 30.5 psi in the ExRPath denaturation buffer to maximize the expansion factor. Such adaptations helped overcome the heavily fixed nature of archival human brain tissue.

To investigate putative interactions of SARS-CoV-2 with the human brain, formalin fixed post-mortem brain tissue of multiple brain regions from both COVID-19 decedents and age-matched non-COVID-19 individuals were prepared for ExRPath, which requires embedding in a polyacrylate hydrogel. The specimen, before hydrogel embedding, had been treated with AcX to equip proteins with hydrogel-binding anchors, and subsequently expanded 4x after protein softening with heat and detergent treatment. The resulting specimens were re-embedded in another round of swellable hydrogel, followed by addition of water, to expand the tissue to 20x of its original size. ExRPath processed tissue was subsequently immunostained (**Fig. 1a**).

**Figure 1:**
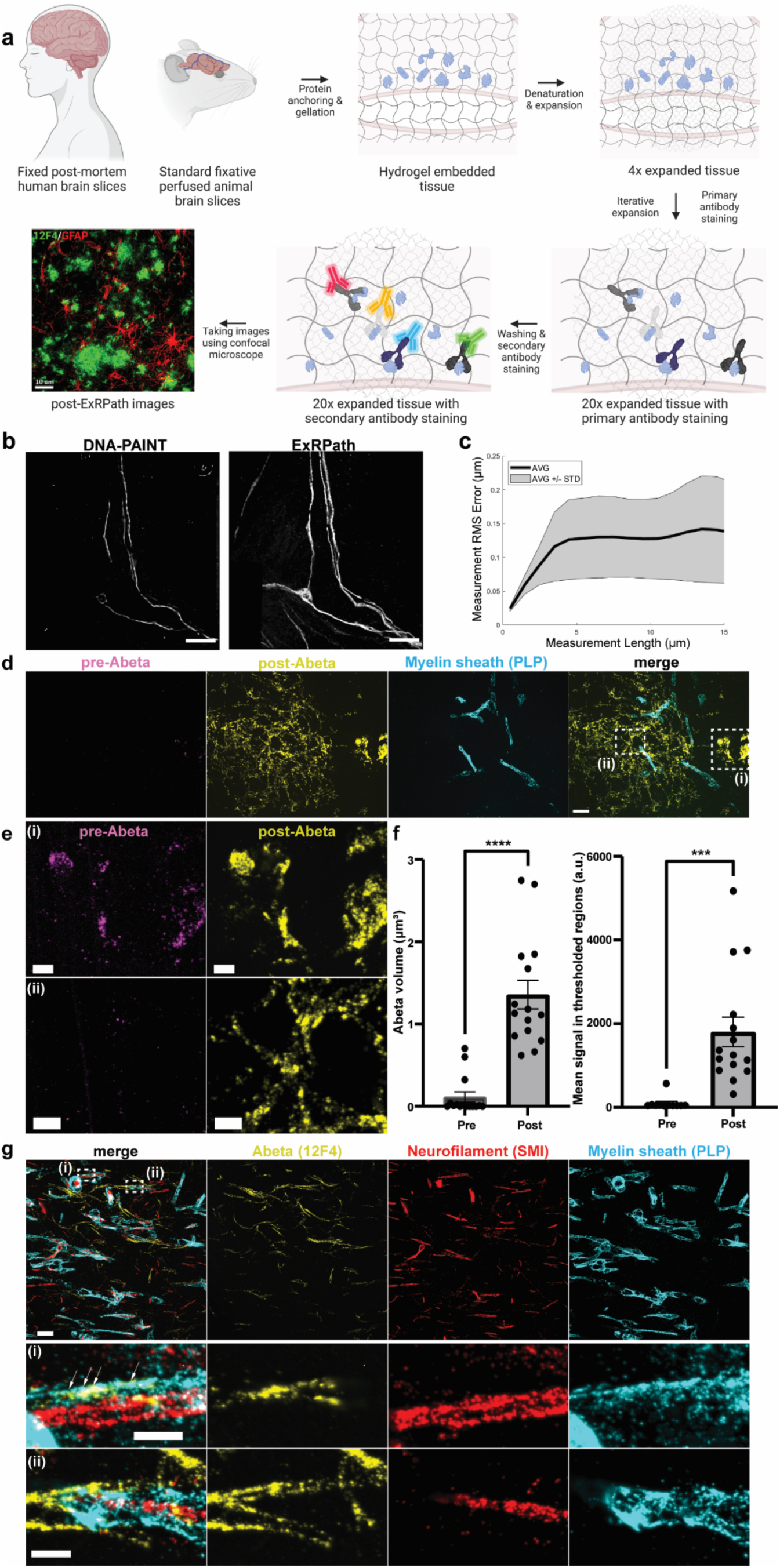
Expansion revealing of pathology (ExRPath) approach. **a)** Schematic overview of ExRPath. Post-mortem human tissue samples (fixed in 10% formalin) or conventionally fixed brain tissue (e.g., formaldehyde preserved mouse brain tissue) are treated with AcX, and embedded in a swellable hydrogel. After denaturation, gels are expanded 4 times in water. After iterative expansion, gels are expanded 20 times, and antibody staining can reveal previously invisible protein nanostructures using conventional microscopes. **b)** Filtered maximum intensity projection of rigid-registered ExRPath image (left, see Methods) and DNA-PAINT rendering (right) of neurofilament (SMI) staining in cultured neurons, obtained from the same field of view (exemplar field of view from 2 fields of view from 1 well from 1 culture batch). **c)** Root mean square error vs measurement length (in biological units), calculated via a non-rigid registration algorithm of DNA-PAINT vs ExRPath-processed cultured neurons (n = 2 fields of view from 1 well from 1 culture batch). Black line, mean; gray shading, area between each field of view and the mean. **d)** ExRPath confocal images (maximum intensity projections) of human COVID+ brain tissue samples showing immunostaining with antibodies against Aβ (12F4) and myelin sheath (PLP). From left to right: merged pre- and post-expansion staining of Aβ42 with post-expansion myelin sheath (PLP), pre-expansion staining of Aβ42 (magenta), post-expansion labelling of Aβ42 (yellow), and post-expansion myelin sheath (cyan) (Scale bar, 2 μm). **e)** Boxed regions in the merged image of panel d. (Scale bar, 500 nm). **f)** Quantification of Aβ showing significantly increased volume (left) and mean intensity (right) in post-ExRPath staining compared to pre-expansion staining (unpaired t-test on difference between pre- and post-expansion staining, n = 15 fields of view from 3 samples from 3 patients, ****p<0.0001, ***p=0.0009), **g)** ExRPath confocal images (maximum intensity projections) showing post-expansion Aβ, neurofilament, and myelin sheath staining (upper, scale bar 2 μm), and **(i-ii)** close-up views of boxed regions from upper panel showing Aβ nanoclusters (bottom, scale bar 500 nm); scale bar of all ExRPath images in biological units (the physical size divided by the expansion factor); shown are images from representative fields of view from two independent replicates (n = 3 tissue sections across n = 3 patients).

To validate ExRPath, we compared post-ExRPath images to pre-expansion DNA-PAINT images of the same cultured neuron sample {Schnizbauer 2017} and found low distortion comparable to that seen from earlier expansion technologies (**Fig. 1b,c**). To quantitatively assess the nanoscale precision of ExRPath, as we did previously for ExR {Sarkar NBME 2022}, we calculated, as a linearized distortion measure between ExRPath and DNA-PAINT, the shifts (in nm) between the half-maxima of the autocorrelation (ExRPath-ExRPath or PAINT-PAINT) and cross-correlation (ExRPath-PAINT) functions for segments of neurofilament imaged using both modalities (**Extended Data Fig. 1a,b**). Our analysis of two regions of interest (ROIs) from one well of cultured neurons showed a mean linearized distortion of 19.14 nm for the ExRPath-PAINT cross-correlation versus PAINT-PAINT autocorrelation (**Extended Data Fig. 1a**, **Extended Data Table 1**; 95% CI of the mean (-28.93, 67.21)) and 15.63 nm for the ExRPath-PAINT cross-correlation versus ExRPath-ExRPath autocorrelation (**Extended Data Fig. 1b**, **Extended Data Table 1**; 95% CI of the mean (-26.66, 57.91)). Thus, the upper bound on total distortion due to ExRPath remains in the low double-digit nanometer range, as with the original ExR technology.

To characterize the decrowding effect of ExRPath, we compared pre-ExRPath to post-ExRPath amyloid-beta (Aβ) staining at the same resolution (**Fig. 1d,e**). The pre-expansion Aβ staining showed signal only in the middle of plaque centers, but the post-expansion staining revealed detailed nanostructures of Aβ, which sometimes colocalized with myelin proteolipid protein (PLP). Quantification of Aβ signal showed significantly increased volume and mean intensity in post-ExRPath staining (**Fig. 1f**). Therefore, just as with ExR, ExRPath can reveal previously invisible protein nanostructures, but now in human brain tissues. Co-staining COVID-19-positive decedent brain tissue for markers of Aβ-42 and myelin revealed periodic Aβ nanoclusters (**Extended Data Fig. 1c,d**) near the myelin sheath (**Fig. 1g**). These images demonstrate the utility of ExRPath in revealing nanoscale pathology signals in human brain tissue samples.

To investigate putative interactions that may contribute to the neurological sequelae associated with SARS-CoV-2 infection, we examined brain tissue of COVID decedents. To identify dysregulated pathways, we performed single-nucleus RNA sequencing (snRNA-seq) on prefrontal cortex (PFC) tissue, an important brain region for cognitive control, executive function, and short-term memory, analyzing samples from 8 COVID-positive decedents compared to transcriptional signatures from an age- and sex-matched control group (**Fig. 2a, Extended Data Table 2**). While infection information is not available, the patients included in the study were deceased prior to February 2021, which is prior to documentation of the spread of SARS-CoV-2 variants in the US and we expect to be of the original strain. The study design aimed to balance age and sex, to the extent possible with the available tissue. The sequenced transcriptomes from each nucleus were clustered via graph-based clustering (UMAP). Cell type identities were assigned to each cluster based on marker gene expression demonstrating coverage across major cell types including oligodendrocytes and their progenitors (OPC), excitatory and inhibitory neurons, microglia, astrocytes, and vascular cells (**Fig. 2b**). COVID-positive decedent tissue had significantly more astrocytes (17.9%) than non-COVID tissue (8.2%), an over two-fold increase (odds ratio of 2.43, p=1.7e-3 by quasi-binomial regression) that was suggestive of a neuroinflammatory phenotype (**Fig. 2c**, **Extended Data Fig. 2a**). An increased number of astrocytes in COVID-positive decedent tissue was similarly observed via histological analysis of tissue specimens for GFAP, an astrocyte marker associated with inflammatory response (**Extended Data Fig. 2b,c**).Cell types were identified in the snRNA-seq data set with canonical markers (**Fig. 2d**) and analyzed for differentially expressed genes (DEGs) (p < 0.05) between the two groups for each of the cell types. Astrocytes displayed significantly increased inflammatory marker glial fibrillary acidic protein (*GFAP*) and microglia-astrocyte crosstalk gene prostaglandin D2 synthase (*PTGDS*). PTGDS is an inflammatory modulator shown to modulate microglial responses, including in the context of a DJ-1 KO injury model, in which rescuing PTGDS expression in astrocytes ameliorated microglia inflammatory signatures” {Choi 2019}. DEGs were dominated by spliceosomal and RNA processing machinery, including *SON* and *PNISR,* and significantly downregulated DEGs associated with synaptic regulation and neuronal regulation like glypican 5 (*GPC5*) and neurexin-1 (*NRXN1*) (**Extended Data Fig. 2d**). Notably, for microglia, the resident immune cell type of the brain, the COVID-positive cohort displayed strong upregulation for a variety of pathways associated with activation, inflammatory signatures, and viral defense responses (**Fig. 2e**). In concordance with a recent snRNA-seq study of choroid plexus and cortex tissues in severe COVID-decedents, we detected an increase in microglia activation genes associated with Alzheimer’s Disease (AD) (e.g., *CD74*, *C1QC*) {Yang 2021}. We also found an increase in other genes associated with MHC Class II (*MHC-II*) protein binding (including *LILRB1*, *CLEC7A*, *HLA-E*, *HLA-DMB*, and *HLA-DRA*) and in genes associated with Aβ binding, including LRP1 and apolipoprotein E (*APOE*), of which allele ε4 is the strongest genetic risk factor for late-onset AD.

**Figure 2:**
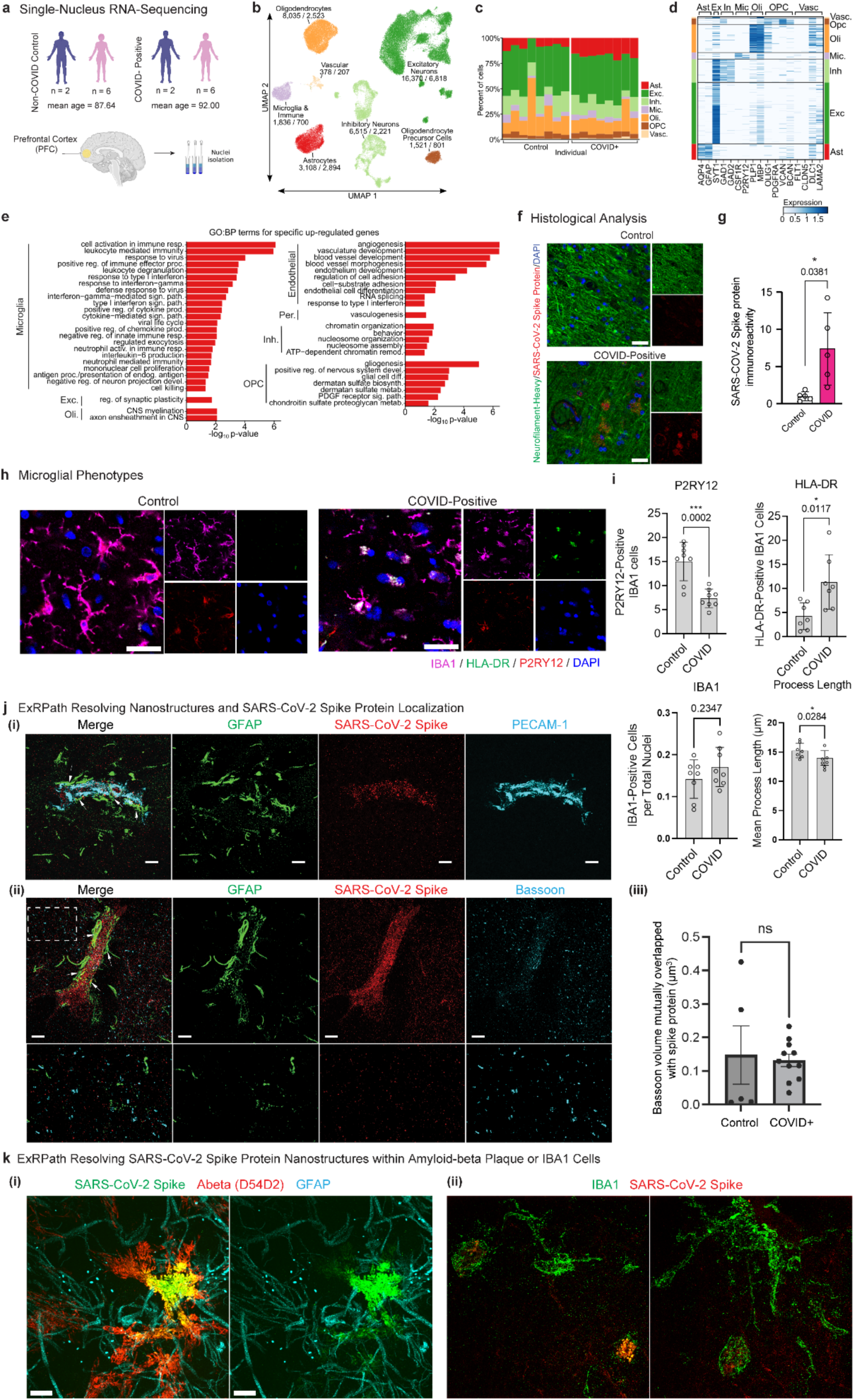
COVID-positive decedent brain tissue displays SARS-CoV-2 immunoreactivity and demonstrates increased inflammatory phenotypes. **a)** COVID-positive and non-COVID control individuals were selected to balance age and sex across groups and nuclei were isolated from flash-frozen samples of PFC, **b)** transcriptional signatures were used to identify cell populations (number of non-COVID/COVID+ nuclei included below each labeled cluster), **c)** ratios between the various cell types for each individual, **d)** canonical cell type-specific genes identifying each cell type (Astrocytes (Ast), Excitatory Neurons (Ex), Inhibitory Neurons (In), Microglia (Mic), Oligodendrocytes (Oli), Oligodendrocyte Precursor Cells (OPC), Vascular Cells (Vasc), **e)** GO:BP terms for upregulated cell type-specific DEGs (1-2 cell types only) for individual cell types, **f)** SARS-CoV-2 immunoreactivity in non-COVID control (left) and COVID-positive decedent (right) PFC tissue (red: SARS-CoV-2 spike protein, green: neurofilament, blue: DAPI; scale bar, 100 µm) characterized via conventional histological and immunohistochemistry methods, **g)** quantification of SARS-COV-2 immunoreactivity (n = 5 images for each of n = 3 tissue sections across n = 3 patients; statistical analysis via t test), **h)** characterization of microglial phenotypes in COVID-positive (left) and non-COVID control (right) PFC tissue (magenta: IBA1, green: HLA-DR, red: P2RY12, blue: DAPI; scale bar, 30 µm) characterized via conventional histological and immunohistochemistry methods, and **i)** quantification between these conditions for the number of P2RY12-positive cells, HLA-DR-positive cells, fraction of IBA1-positive cells determined by positive cells out of total number of nuclei, and average length of microglia processes (n = 8 tissue sections for P2RY12 and IBA1, n = 7 tissue sections for HLA-DR and process length, each across n = 3 patients per group; statistical analysis via t test), **j)** ExRPath confocal images of max intensity projection after immunostaining with antibodies against **(i)** GFAP, SARS-CoV-2, and PECAM-1, and **(ii)** GFAP, SARS-CoV-2, and bassoon. Arrows indicated visualization via ExRPATH of SARS-CoV-2 localization around astrocytes in COVID-19 decedent tissue (scale bar, 2 µm), **(iii)** volume mutually overlapped between bassoon and spike protein (p = 0.7965, unpaired t-test on bassoon-spike overlap, n = 11 fields of view from 2 human COVID decedent tissue specimens, n = 5 fields of view from 2 human control specimens). **k)** ExRPath confocal images of max intensity projection after immunostaining with antibodies against **(i)** GFAP, SARS-CoV-2, and D54D2, and **(ii)** SARS-CoV-2 and Iba1 (scale bar, 2 µm). Shown are 2 human COVID decedent tissue specimens selected from 8 patients with multiple brain tissues analyzed in a rare minority patient specimen.

Vascular cell types displayed enhanced angiogenic and vascular development pathways and excitatory neurons displayed increased expression of genes associated with the regulation of synaptic plasticity (**Fig. 2e**). Comparing cell types, a subset of DEGs were shared across multiple cell types, in addition to cell type-specific DEGs (**Extended Data Fig. 2e**), where 219 DEGs were shared by at least 3 cell types and 32 DEGs were shared by at least 7 cell types (**Extended Data Fig. 2f**), and these shared DEGs were expressed across cell types, without any one cell type expressing significantly more shared DEGs (**Extended Data Fig. 2g**). Among DEGs shared by 3 or more cell types, genes associated with mRNA processing were among those most upregulated in the COVID-positive group, while those associated with cell junction and neuron morphology, connectivity, and organization were among those most significantly downregulated (**Extended Data Fig. 2h**).

As the transcriptomic analysis evidenced prominent changes in mRNA processing and metabolic processes, we sought to further profile the presence and localization of SARS-CoV-2 viral elements in the brain. We detected immunoreactivity to the SARS-CoV-2 spike protein in the PFC (**Fig. 2f,g**) as well as in other brain regions examined, including the olfactory bulb {Meinhardt 2021} (**Extended Data Fig. 3a,b**). Assessing the number of IBA1-positive microglia in the PFC from COVID and non-COVID decedents, we did not detect significant differences in numbers (p = 0.2347). Co-staining brain tissue sections for the microglial homeostatic marker purinergic receptor (P2RY12) and IBA-1, we found a significant (p = 0.0002) decrease in the number of IBA1-positive microglia co-expressing P2RY12 in the PFC of COVID-positive decedent (**Fig. 2h,i**). Concordantly, the length of processes in microglial of COVID-positive PFC were significantly (p = 0.0284) decreased compared to non-COVID PFC, indicative of an increase in motile, inflammatory cells (**Fig. 2h,i**). Notably, the MHC-II receptor and T-cell activation marker HLA-DR, which have been associated with AD {Bryan 2008; Swanson 2020; Mathys 2017}, was significantly (p = 0.0117) increased in COVID-positive tissue. Collectively, this is consistent with the interpretation that microglia in the COVID-decedent brain are in a neuroinflammatory state.

To gain insights into putative interactions between SARS-CoV-2 proteins and brain proteins, we applied ExRPath to tissues in which we had detected viral spike protein (**Fig. 2j**). We analyzed brain tissue from 8 COVID-19 patients and 8 control individuals. Spike protein immunoreactivity was detected in 2 out of 8 COVID-19 patient specimens. Both of these specimens exhibited amyloid deposits, with approximately 10% of spike signal overlapping with amyloid. Among the remaining 6 COVID-19 specimens, no spike signal was detected, although all contained amyloid deposits. In the 8 control specimens, spike protein was not detected and amyloid signal was comparatively lower. This highlights the utility of ExRPath as a high-throughput method for surveying large volumes of human brain tissue to identify rare molecular overlaps.

Given the increased inflammatory phenotypes revealed via snRNA-seq, we sought to further investigate the association of SARS-CoV-2 proteins with inflammatory cell types. ExRPath revealed SARS-CoV-2 spike protein immunoreactivity in close registration with astrocytic processes (**Fig. 2j**), which at higher resolution could be localized in vessel tubule structures positive for PECAM-1 (platelet endothelial cell adhesion molecule) and surrounded by astrocytic endfeet (**Fig. 2j**). To further probe viral protein element localization, we investigated whether SARS-CoV-2 spike proteins colocalize with synapses. Co-staining spike protein with presynaptic protein bassoon, we observed no such colocalization (**Fig. 2jii**). No difference in overlapping signal was found between COVID-positive and non-COVID control tissue samples (**Fig. 2jiii**). As evidence suggests Aβ is associated with antiviral immune responses in the brain {Wozniak 2007; Kumar 2016; Eimer 2018}, we further examined spike-positive tissues, co-labeling Aβ and SARS-CoV-2 spike protein via ExRPath. While both Aβ and spike viral protein elements are expected to be low in abundance in the human brain, we did detect overlapping signal between Aβ and SARS-CoV-2 spike protein (**Fig. 2ki, Extended Data Fig. 1ei**). To quantify the colocalization of Aβ and SARS-CoV-2 spike protein at the nanoscale, we used Aβ punctae as reference points and extended a search radius outward. We measured the mean number of spike punctae within each radius away from an Aβ puncta. At all search radii examined, the observed number of colocalized punctae was significantly greater than what would be expected if punctae were randomly distributed, which demonstrates non-random colocalization of SARS-CoV-2 viral proteins and amyloid, and potentially pointing towards a biologically relevant interaction (**Extended Data Fig. 1eii-iii**). Furthermore, to explore the relationship between viral proteins and immune activation, we co-labeled IBA-1 (a marker of microglia) and SARS-CoV-2 spike protein. ExRPath imaging revealed engulfed spike proteins within IBA-1–positive microglial cell bodies, suggesting active involvement of microglia in viral protein processing (**Fig. 2kii**). While we here focused the spike protein, further work could profile the relative abundance of the nucleocapsid protein and correlates of direct viral infection.

To further evaluate the functional relevance of a SARS-CoV-2 spike protein and Aβ combination in a model inclusive of amyloid plaques, with which to probe localization and immune response, we sterotactically administered S1 spike protein intracranially to the cortex of mice. As wild-type mice exhibit low levels of brain amyloid, we employed the well-established 5XFAD amyloid mouse model that begins to develop amyloid plaques by age two to four months {Oakley 2006i}. We stereotactically injected SARS-CoV-2 S1 viral spike protein or vehicle control intracranially into the cortex (-1.8 mm anteroposterior, 2.2 mm lateral, relative to bregma) of four month-old 5XFAD mice (**Fig. 3a**). Mice were sacrificed 2 weeks later and microglia phenotypes and amyloid plaques were analyzed. While this approach is less physiologically relevant than administration of live virus, it has enabled protein co-localization and sequelae analyses while requiring less biosafety precautions, and appropriate given that the goal of this technical report is to show the kinds of data that can be generated with our technology.

**Figure 3:**
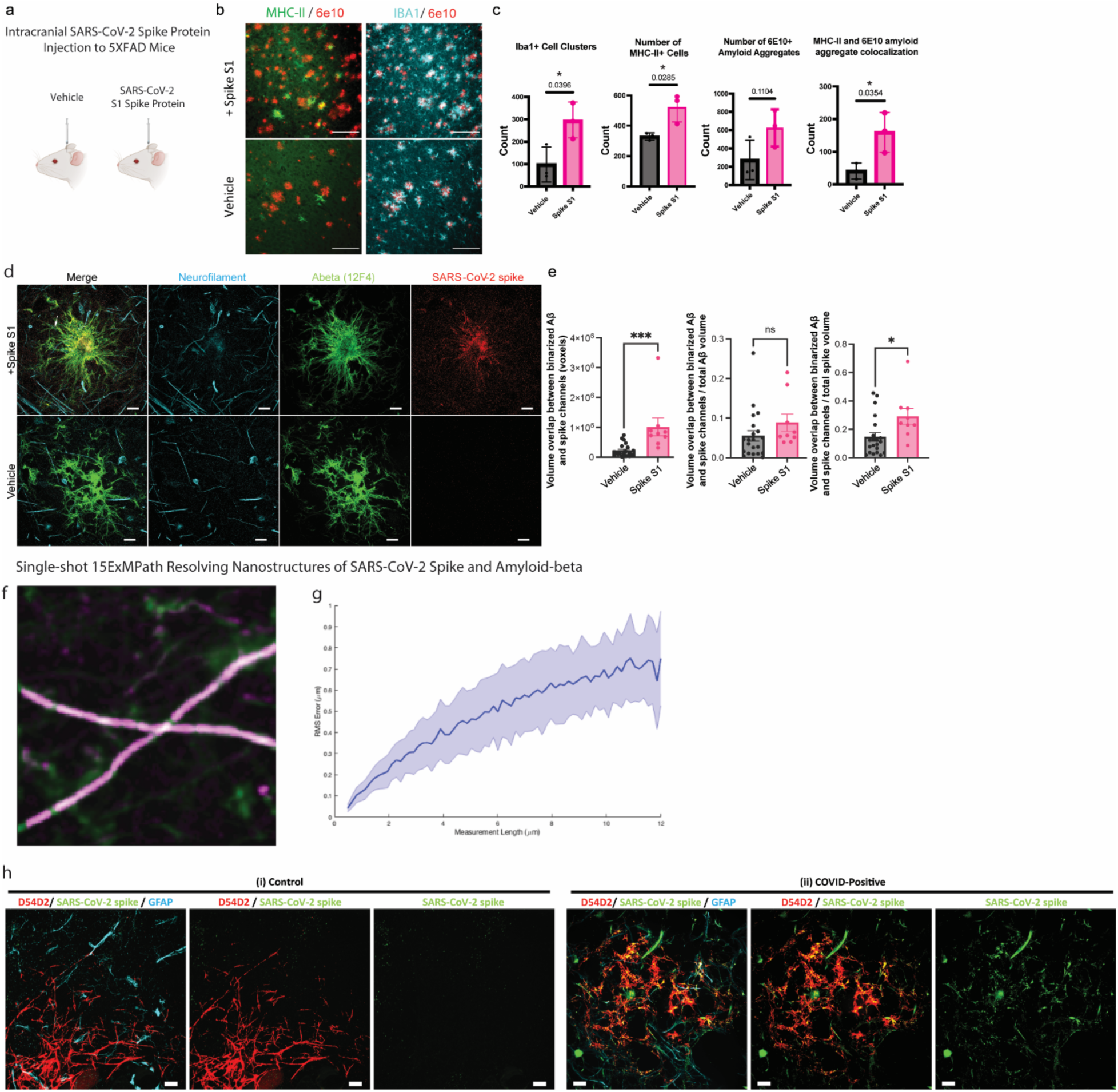
Combination of amyloid and SARS-CoV-2 spike protein induces distinct inflammatory phenotype. **a)** Schematic of intracranial administration to 5XFAD mouse model of SARS-CoV-2 spike protein or vehicle control, **b)** mouse brain sections co-stained for Aβ (red: 6e10) and AD-associated microglia marker MHC-II and pan-microglia marker IBA1 (green: MHC-II, cyan: IBA1; scale bar, 100 µm), **c)** quantification of (left to right) the number of IBA1-positive cell clusters, the number of MHC-II-positive cells, the colocalization of MHC-II and 6e10, and the number of Aβ aggregates (n = 3 mice per group, data points representing averages for each animal; quantification based on 150 cells for IBA1 and MHC-II, 275 D54D2-positive regions, and 275 6e10-positive regions; statistical significance determined by unpaired t-test), **d)** ExRPath confocal images (maximum projection intensity) of (top) tissue with SARS-CoV-2 spike protein injected compared to (bottom) vehicle controls for SARS-CoV-2 and amyloid immunoreactivity (cyan: SMI neurofilament, green: 12F4 amyloid, red: SARS-CoV-2; scale bars, 2 µm); shown are images from representative fields of view from two independent replicates (3 mice per group), and **e)** quantification of SARS-CoV-2 and amyloid co-localization in 5XFAD mouse tissue samples in terms of (left) volume of mutually overlapping signal (see Methods), (middle) volume of overlap normalized by the amyloid plaque volume, (right) volume of overlap normalized by spike volume (n = 20 fields of view in the vehicle group and n=8-9 fields of view in the Spike S1 group from 3 mice per group; statistical significance as determined by an unpaired t-test, *p<0.05, ****p<0.0001, p < 0.0001 for mutual volume overlap; p = 0.1938 for mutual volume overlap normalized to amyloid volume; p = 0.0198 for mutual volume overlap normalized to spike protein volume). **f)** Nonrigidly registered pre-expansion x40 magnification confocal image (green) and post-expansion x4 magnification confocal image (magenta) of the same control human tissues (from one representative experiment of one brain slices from two individuals) with 15ExMPath protocol. **g)** RMA measurement error as a function of measurement length of data acquired as in f (blue line, mean; shaded area, ±1 s.d.; n=6 areas from two brain slices from two individuals). **h)** Single-shot 15ExMPath confocal images (maximum projection intensity) of (i) control tissue and (ii) COVID-positive tissue after immunostaining with antibodies against (i) GFAP, SARS-CoV-2, and D54D2 (scale bars, 2 µm). Shown are representative fields of view from 3 independent replicates across 8 patient specimens, with multiple brain sections analyzed (3 cases per group).

5XFAD mice injected with S1 spike exhibited a significantly increased clustering of microglia compared to vehicle injected mice, determined by the number of regions (defined as 15 μm regions using Imaris) containing multiple IBA1-positive cells (**Fig. 3b,c**). Consistent with an inflammatory response, S1 spike protein-injected 5XFAD mice also exhibited significantly (p = 0.0396 and 0.0285) increased numbers of MHC-II-positive cells compared to vehicle injected mice suggesting that S1 spike protein increases inflammatory microglial response in 5XFAD mice (**Fig. 3b,c**). To determine whether the increase of MHC-II colocalizes with Aβ aggregates, we quantified the number of colocalized MHC-II-positive cells and Aβ-positive regions. While S1 spike injected mice tissue did not exhibit a significant increase in Aβ immunoreactivity, the Aβ-positive regions colocalized with MHC-II-positive cells were significantly increased (p = 0.0354) compared to vehicle injected 5XFAD mice. This suggests that the combination of SARS-CoV-2 spike protein and amyloid may trigger an inflammatory microglia response *in vivo*. (**Fig. 3b,c**).

We thus sought to visualize SARS-CoV-2 S1 spike protein localization in mouse brain tissue with high resolution. Applying ExRPath strikingly revealed SARS-CoV-2 spike protein colocalization within Aβ plaques, with the spike protein detected throughout the Aβ plaques and at a high density at the plaque cores (**Fig. 3d,e**). The overlap between amyloid and spike signals, quantified as the fraction of total volume of one channel mutually overlapping with the other, was significantly increased in spike protein-administered animals compared to controls (**Fig. 3e**). Normalizing this volume overlap by total spike signal volume resulted in a statistically significant increase, while normalizing by total amyloid volume did not. That is, the relative proportion of spike volume co-localized with amyloid, but not the proportion of amyloid volume co-localized with spike, was higher in 5XFAD mice injected with spike protein, suggestive of an amyloid-spike interaction dictated by the amount of spike protein. Thus, following intercranial injection of SARS-CoV-2 Spike S1 protein, 5XFAD mice display a colocalization between Aβ and spike protein and an increase in neuroinflammatory phenotypes, with AD-associated signatures.

We recently developed single-shot 20-fold expansion microscopy (20ExM), which achieves ∼20-nm resolution on conventional microscopes through a single 20× physical expansion step, validated on mouse brain tissue and cultured neurons {Wang, 2024, 20ExM}. Since 20ExM supports post-expansion immunostaining in mouse brain tissue, we here optimized the protocol for post-mortem human brain specimens, resulting in a single-shot 15x expansion protocol for human brain tissue, which we here designated 15ExMPath. The adaptation process involved modifying the protein anchoring and denaturation procedures described above. The expansion factor obtained with 15ExMPath is somewhat lower than that achieved in mouse tissue (20x), perhaps reflecting the effects of long-term fixation on human specimens. We tested EDTA and beta-mercaptoethanol treatments to improve fixed-tissue softening, but these modifications did not increase expansion to the full 20x observed in mouse tissue, suggesting that very large expansions of human tissue in single steps might require qualitatively different softening methods from what has been used before. We validated the macroscopic distortion introduced by 15ExMPath using established ExM distortion analysis. This approach calculates the root mean square (RMS) alignment error from the deformation vector field generated by registering images of the same field of view before and after expansion. To perform this analysis, we applied 15ExMPath to formalin-fixed human control brain slices, followed by post-expansion staining against neurofilament proteins **(Fig. 3f)**. The resulting RMS error was ∼6%, comparable to values reported for previous post-expansion staining ExM protocols for human tissue **(Fig. 3g)**. Using this optimized approach, we performed immunostaining for Aβ (D54D2), SARS-CoV-2 spike protein, and GFAP, and again observed strong spatial co-localization between Aβ and spike proteins **(Fig. 3h)**, as we saw with ExRPath.

Each of ExRPath and 15ExMPath has distinct advantages and limitations. ExRPath involves more hands-on steps – 1^st^ round swellable gelling, 2^nd^ round re-embedding, 3^rd^ round swellable gelling - but the entire protocol can be performed on an open bench. Gel handling is straightforward due to robust gel integrity, and the final expansion factor is reliably ∼20x. In contrast, 15ExMPath requires only a single gelling step, reducing hands-on time while still achieving a high expansion factor. The procedure must be conducted in an inert nitrogen environment (i.e., in a glove bag), with precise gelation termination, to achieve the final expansion factor and preserve gel integrity. Nevertheless, 15ExMPath achieves the highest single-step expansion factor reported for human tissue samples among all published expansion microscopy protocols. Taken together, users should select the appropriate protocol based on their experimental goals and available resources.

Together, these results suggest that SARS-CoV-2 spike proteins induce neuroinflammatory responses and that the combined presence of spike proteins and Aβ might induce unique inflammatory responses. We have developed novel techniques, ExRPath and 15ExMPath, to expand standard human pathology brain specimens to visualize nanoscale, hidden protein localization. Coupling ExRPath technology to a mouse model and leveraging iPSC-based *in vitro* approaches, we further identified CNS pathways and phenotypes perturbed by the SARS-CoV-2 virus in human decedent brain tissue. In each of these model systems, the SARS-CoV-2 spike protein induces inflammatory phenotypes in microglia. While the young, healthy human brain contains low levels of Aβ and the pathological samples we analyzed contain only low levels of viral protein elements, we did observe colocalization between Aβ and SARS-CoV-2 spike proteins and identified distinct microglial inflammatory signatures that can result from co-administration of these species *in vitro.* In the Aβ-abundant 5XFAD mouse model, these spike proteins colocalize with Aβ plaques and exacerbate microglia inflammation. These data motivate further work to understand the downstream sequelae of this acute effect, the duration and temporal resolution of the inflammatory signatures, and any potential persistent or long-term effects. A recent study probing the association between viral exposure and neurodegenerative disease found that some viral exposures increased risk of neurodegeneration up to 15 years from the time of infection {Levine 2023}. This prompts the question: what is the risk profile associated with SARS-CoV-2? Further motivation for investigation into long-term sequelae is garnered from the prevalence of patients suffering from long-COVID and from persistent neurological symptoms after recovery. Collectively, this study demonstrates that ExRPath and 15ExMPath technologies enable nanoscale investigation of disease mechanisms in human pathology samples, a critical step toward understanding disease pathogenesis and identifying new therapeutic opportunities.

## Acknowledgements

L.-H.T. acknowledges funding support of NIH 3-UG3-NS115064-01S1, The JBP Foundation, Cure Alzheimer’s Fund, Lester A. Gimpelson, and Jay L and Carol D Miller . E.S.B. acknowledges the support of Tom Stocky, NIH 1R01EB024261, Kathleen Octavio, Good Ventures, Lisa Yang, NIH 1R01AG070831, HHMI, NIH 1R01MH123403, European Research Council (ERC) under the European Union’s Horizon 2020 research and innovation programme (grant agreement No 835102), NIH 1R56AG069192, NIH R01MH124606, and John Doerr.

A.E.S. was also supported by 1F32AG072813-01. J.W.B was also supported R01NS14239, Cure Alzheimer’s Fund, NASA 80ARCO22CA004, Chan-Zuckerberg Initiative, MJFF/ASAP Foundation, and Brain Injury Association of America. M.E.S. was supported by the NSF Graduate Research Fellowship Program and the MathWorks Fellowship. ROSMAP is supported by NIA grants P30AG10161, P30AG72975, R01AG15819, R01AG17917, and R01AG058639.

This work was performed in part in the Ragon Institute BSL3 core, which is supported by the NIH-funded Harvard University Center for AIDS Research (P30 AI060354).

## Methods

### Cultured-neuron preparation

All experiments were performed according to the Guide for the Care and Use of Laboratory Animals and were approved by the National Institutes of Health and the Committee on Animal Care at the Massachusetts Institute of Technology. Cultured mouse hippocampal neurons were prepared from postnatal ∼day 0 or 1 Swiss Webster mice (Taconic) as previously described {Sakar, 2022}. In summary, 5,000 – 10,000 cells were plated in each well of diluted Matrigel (250 μL Matrigel in 12 mL DMEM (Dulbecco’s Modified Eagle Medium)) treated imaging chambers (Grace Bio Lab, CultureWell^TM^ removable chambered coverglass), and grown at 37°C and 5% CO_2_ in a humidified atmosphere for 14 days before fixation. The cells were briefly washed with PBS, and fixed with 4% paraformaldehyde for 6 hrs at 37°C. Then, samples were washed with PBS after removing fixative, and stored at 4°C.

## Human brain tissue preparation

The study of human tissue specimens was approved by the Massachusetts Institute of Technology’s Committee on the Use of Humans as Experimental Subjects. Brain tissue samples came from participants enrolled in the Religious Order Study (ROS) or the Rush Memory and Aging Project (MAP), longitudinal studies of aging and dementia, collectively known as ROSMAP. Age- and sex-matched groups of COVID-positive and non-COVID decedents were selected, with 2 males and 6 females in each group and with mean ages of 92.00 and 87.64, respectively. Decedent brain tissue samples were prepared as fresh-frozen samples for snRNA-seq and fixed samples for histology. Samples fixed in 10% formalin for 1 month were sectioned with a microvibratome into 20 μm slices, and stored in PBS at 4°C.

## Mouse brain tissue preparation

All experiments were performed according to the Guide for the Care and Use of Laboratory Animals and were approved by the National Institutes of Health and the Committee on Animal Care at the Massachusetts Institute of Technology. 5XFAD mice were obtained from Jackson Laboratory (JAX ID# 34840). A total of 10, 4-month-old 5XFAD female mice were used for these studies. Mice were housed in groups of five animals on a standard 12 h light/12 h dark cycle, temperature (24 °C), and humidity (45%) with food and water available ad libitum. All experiments were performed during the light cycle. After SARS-CoV-2 spike protein injection, mice were perfused transcardially with ice-cold 4% formaldehyde solution (Fisher Scientific) in sodium phosphate buffer. Brains were post-fixed with 4% formaldehyde solution for 12 hours, before being sectioned at 50 μm using a vibratome (Leica).

## ExRPath of human and mouse tissue slices and cultured neurons

Acryloyl-X, SE (Life Technologies, A20770) (AcX) stock solution was prepared by adding anhydrous dimethylsulfoxide (DMSO) to AcX powder for a 10 mg/mL concentration. Brain slices or cultured neurons were treated with diluted AcX solution (0.1 mg/mL) in PBST (1x PBS with 0.1 % Triton X-100) and incubated for 2 hrs at 37°C. Samples were washed 3 times using PBST for 5 min each at room temperature and placed on a cover glass for gelation. For slice samples, gel chambers were assembled between cover glasses with spacers of two #1.5 cover glasses. The first gelling solution was injected between cover glasses to cover slice samples entirely. For cultured neurons, the first gelling solution was deposited into the imaging chamber fully (50 μL), and each well was covered on top using a cover glass. Samples were incubated at 4°C for 30 min, then 37°C for 30 min. Gels containing brain slices or cultured neurons were cut out from surrounding excess gel and incubated for 60 min in a benchtop autoclave (VWR, CLS-1191) at 121°C and 30.5 psi in denaturation buffer (see Table below for all compositions) as previously described {Valdes, 2021, dExPath} with slight modification. To be specific, β-mercaptoethanol was excluded to preserve pre-expansion stained antibodies for comparison between pre- vs. post-expansion staining efficiency, and we found that β-mercaptoethanol was not needed for isotropic expansion or final expansion factor. We also found that a lower concentration of SDS than the dExPath protocol was easier to use, to prepare the denaturation buffer, when using solution-type SDS stock solutions. After denaturation, gels were washed 3 times with excess water for 15 min each.

For re-embedding, expanded first gels were incubated with re-embedding solution on a shaker at room temperature twice for 1 hr each. Gels were gently scooped up using disposable spatulas (VWR, 80081-192), and placed between cover glasses. After nitrogen purging in a ziploc bag, gels were incubated at 45°C for 1-2 hrs. The re-embedded gels were incubated with a 3rd gelling solution on a shaker at room temperature twice for 1 hr each. The gels were placed between cover glasses without bubbles by adding access gelling solution and incubated at 60°C for 1 hr. Gels were washed several times with excess water and trimmed axially to 1 mm thickness to facilitate subsequent immunostaining and imaging of gels.

Gelling solution composition Monomer solution:

| Component | Stock Concentration* | Amount (mL) |
| --- | --- | --- |
| Sodium acrylate | 38 | 9 |
| Acrylamide | 50 | 2 |
| Sodium Chloride | 29.2 (5 M) | 16 |
| PBS | 10x | 4 |
| Water | - | 3.6 |
| Total | - | 34.6 |

\* All concentrations in g/100 mL except PBS

Gelling solution of ExRPath:

| Reagent | Stock Concentration | 1<sup>st</sup> gel solution (μL) | Re-embedding solution (μL) | 3<sup>rd</sup> gel solution (μL) |
| --- | --- | --- | --- | --- |
| Monomer | - | 864 | - | 864 |
| Acrylamide | 50 | - | 275 | - |
| Bis acrylamide | 2 | 40 | 18.75 | 20 |
| Water | - | 36 | 701.25 | 111 |
| 4HT | 0.5 | 20 | - | - |
| TEMED | 10 | 20 | 2.5 | 2.5 |
| APS | 10 | 20 | 2.5 | 2.5 |
| Total (mL) | - | 1 | 1 | 1 |

Denaturation buffer:

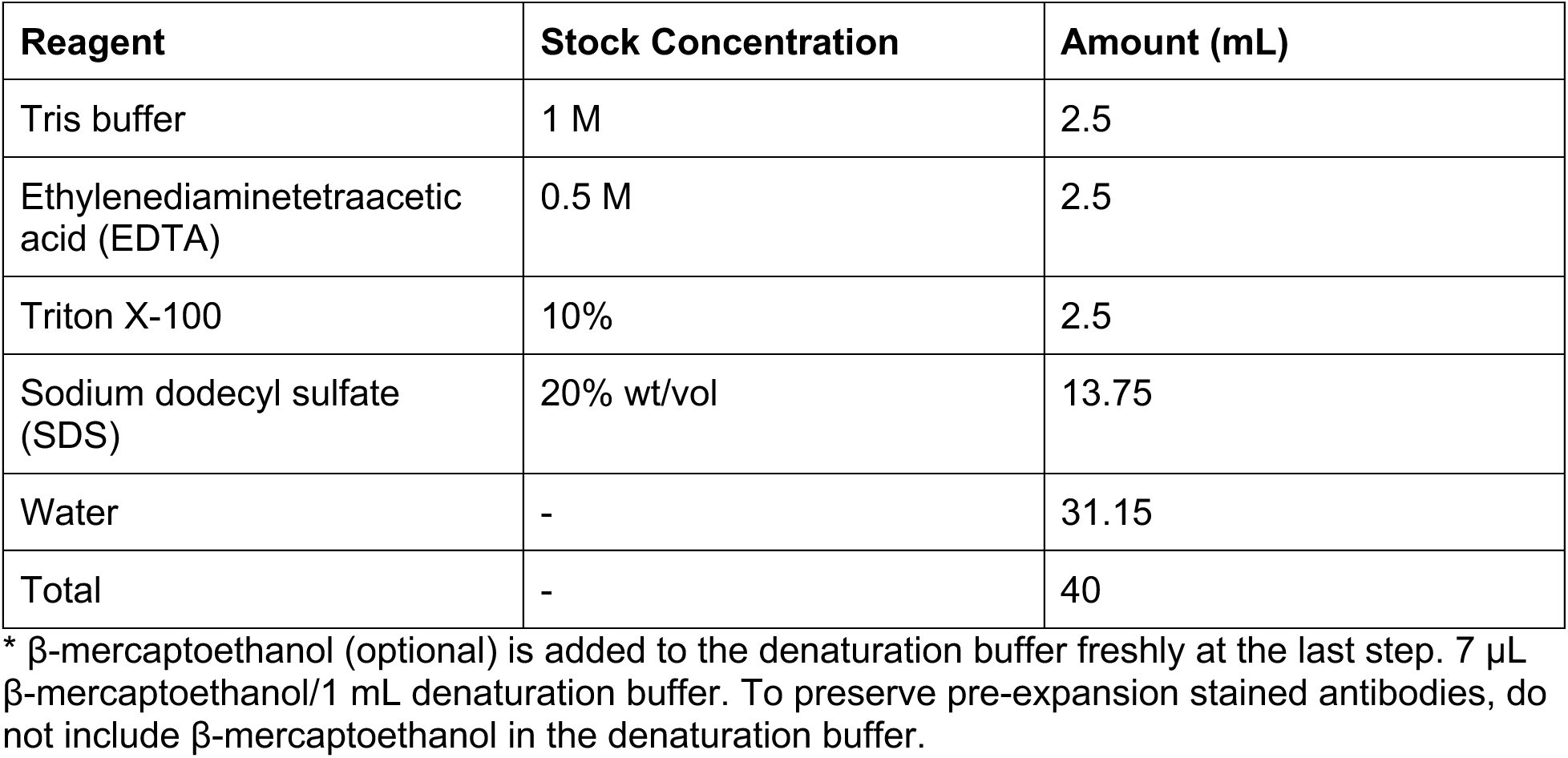

| Reagent | Stock Concentration | Amount (mL) |
| --- | --- | --- |
| Tris buffer | 1 M | 2.5 |
| Ethylenediaminetetraacetic acid (EDTA) | 0.5 M | 2.5 |
| Triton X-100 | 10% | 2.5 |
| Sodium dodecyl sulfate (SDS) | 20% wt/vol | 13.75 |
| Water | - | 31.15 |
| Total | - | 40 |

\* β-mercaptoethanol (optional) is added to the denaturation buffer freshly at the last step. 7 μL β-mercaptoethanol/1 mL denaturation buffer. To preserve pre-expansion stained antibodies, do not include β-mercaptoethanol in the denaturation buffer.

Immunostaining of ExRPath samples

Samples were blocked using blocking buffer (0.5% Triton X-100, 5% normal donkey serum (NDS) in PBS) for 1-2 hrs on a shaker at room temperature. Gels were incubated with primary antibodies in the staining buffer (0.25% Triton X-100, 5% NDS in PBS) overnight at 4°C and washed using 1x PBST (1x PBS, 0.1% Triton X-100) 6 times for 30 min each. Gels were incubated with secondary antibodies in staining buffer overnight at 4°C and washed using 0.05x PBST (20x diluted 1x PBS, 0.1% Triton X-100) 6 times for 30 min each. ExRPath samples were imaged using a Nikon CSU-SoRa confocal microscope with 100% laser power and 1 sec exposure time per channel.

## Decrowding experiments to compare pre- vs. post-ExRPath staining

For pre-expansion antibody staining (Fig. 1d), primary antibodies against 12F4 were added to brain slices in staining buffer overnight at 4°C. Stained slices were washed using 1x PBST six times for 15 min each on a shaker at room temperature. Secondary antibodies (goat anti-mouse, 50 μL of 2 mg/mL) were incubated with AcX (2 μL of 1 mg/mL) overnight at room temperature to prepare AcX-conjugated secondary antibodies. Primary antibody-stained slices were incubated with AcX-secondary antibodies in staining buffer overnight at 4°C, and washed using 1x PBST six times for 15 min each at room temperature. Tissue expansion was carried out as previously described without β-mercaptoethanol to maintain AcX-conjugated secondary antibodies in the hydrogel. The tertiary antibodies (donkey anti-goat) were stained after expansion to bind against AcX-secondary antibodies to visualize pre-expansion staining. For post-expansion antibody staining, primary and secondary antibodies without AcX conjugation were added to expanded gels.

## ExRPath image processing and quantification

First, background was subtracted from image stacks using ImageJ/Fiji’s Rolling Ball algorithm with a radius of 50 pixels. To quantify the volume and signal intensity of Aβ in pre- and post-expansion stained ExRPath images (**Fig. 1F**), images were binarized using an intensity threshold set at 4 standard deviations above the mean pixel intensity for the entire image volume. Thresholded volumes were then passed through a 3D median filter of radius 5x5x3 pixels and subjected to size filtration to remove objects smaller than 100 voxels and larger than 5,000 voxels. Volume was calculated as the number of nonzero pixels in the thresholded, filtered image, and mean signal was calculated as the mean pixel intensity value in nonzero pixels.

To quantify the colocalization of Aβ, bassoon, and spike protein (**Fig. 2jiii**, **Fig. 3J**), we automatically identified and quantified the volume of Aβ and spike puncta. Binary image segmentations were created in CellProfiler 4.0 {Stirling 2021}. Segmentations for each channel were created as follows: intensity rescaling (min-max normalize), intensity thresholding (minimum cross-entropy, default parameters with an empirically-determined threshold correction factor), and watershed segmentation (footprint = 10, downsample = 4). We then used MATLAB to apply additional median and size filtration steps and calculate the volume overlapped between Aβ and spike protein segmentations (calculated as the number of nonzero pixels in the intersection between the binarized image volumes of the two channels, either un-normalized or divided by the number of nonzero pixels in the segmentation of beta-amyloid or spike segmentations). In post-expansion units, 1 voxel is equivalent to ∼9x9x22nm^3^.

To quantify the periodicity of the Aβ clusters along the axons (**Extended Data Fig. 1c,d**), we automatically identified and quantified the volume of Aβ. We identified the local maximum of each Aβ cluster with the peak_local_max function in the Python package skimage. We then calculated the distances between adjacent local maximum as a measure of distance between adjacent puncta. We then plotted the histogram showing the distribution of such distances with a total of 20 bins.

To quantify the colocalization of Aβ and spike protein (**Extended Data Fig. 1eii-iii**), we represented the positions of Aβ puncta (blue) and spike protein (orange) with their local maximum in the 3D volume We determined the number of COVID spike protein puncta that overlapped with seed beta-amyloid puncta within a defined radius and compared it to the number obtained from shuffling the COVID spike proteins. A one tailed t-test was performed on the observed and randomized counts at each radius to determine if the observed count is significantly greater than the randomized group.

## Comparison between ExRPath and DNA-PAINT

### Antibody conjugation for DNA-PAINT

The antibody was conjugated with DNA via DBCO-azide click reaction. Briefly, 100 μg carrier-free antibody (without BSA or sodium azide) formulated in 1× PBS was washed and concentrated using Amicon Ultra Filters (50kDa, EMDMillipore, UFC505096) via three rounds of centrifugation at 8000g for 6 mins. Then, fresh-prepared crosslinker (DBCO-sulfo-NHS ester, dissolved in DMF) was added to antibody in the same Amicon filter at a 5:1 molar ratio (crosslinker : antibody = 5:1) and reacted for 1 hour at 4°C. Next, excess crosslinker was removed by three rounds of centrifugation at 8000g for 6 mins. Azide-modified DNA was then mixed with antibody in the same Amicon filter at a 5:1 molar ratio (DNA: antibody = 5:1) and reacted overnight at 4 °C. Final conjugated antibodies were washed five times with 1× PBS in Amicon Ultra Filters at 8000g for 6 mins to remove unreacted DNA oligonucleotides. The concentration of conjugated antibody was measured using Qubit™ protein assay (ThermoFisher, Q33211) and antibody was kept at 4 °C until use.

### Antibodies for DNA-PAINT

Antibodies for DNA-PAINT:
| Target | Antibody | Species | Dilution |
| --- | --- | --- | --- |
| SMI | Biolegend 837904 | mouse | 10 µg/ml |
| Anti-mouse | Jackson ImmunoResearch Laboratories 715-005-151 | donkey | 20 µg/ml |

### DNA sequences (5’ to 3’)

DNA sequences (5' to 3'):
| ID | Sequence | Vendor |
| --- | --- | --- |
| Docking | /5AzideN/ATACATCT | Integrated DNA Technologies |
| Imager | AGATGTAT /Atto-655/ | Biosynthesis |

### Immunostaining for DNA-PAINT

Neuron cultures were blocked and permeabilized in blocking solution (0.3% Triton X-100, 5% IgG-free BS, 1× PBS) for one hour at room temperature. Cell cultures were incubated with primary antibodies in incubation solution (0.2% Triton X-100, 3% IgG-free BSA, 1× PBS) overnight at 4 °C and washed with washing solution (0.3% Triton X-100, 1× PBS) five times (10 mins each). Both dye-labeled and DNA-conjugated secondary antibodies were diluted in incubation solution and incubated with samples for 2 hours at room temperature and then washed with washing solution five times. 1:100 dilution of 100 nm gold nanoparticles (753688-25ML, Sigma-Aldrich) was then added to wells and gently spun down in a centrifuge (500g, 5 mins). Finally, the cells were briefly washed three times with 1× PBS+500 mM NaCl.

### Super-resolution imaging (DNA-PAINT)

Super-resolution imaging was accomplished on a Nikon Eclipse Ti microscope which was operated with (1) Nikon Elements software, (2) 1.49 NA CFI Apo 100x objective, (3) perfect focus system, and (4) total internal reflection fluorescence (TIRF) laser (488nm, 561nm, 647 nm). Lasers were operated in TIRF mode for all acquisitions. For image acquisition, an electron multiplying charge coupled device (iXon X3 DU-897, Andor Technologies) was used. Cameras were operated at 5-MHz refresh rate with 100 EM gain and a 50 ms exposure time. A 1.5x lens was introduced into the optical path allowed for imaging with a pixel size of 110 nm. Images were acquired using RAM capture via a Nikon Fast Timelapse acquisition. 10000-20000 frames of single molecule image stacks were then acquired and analyzed in Picasso. To correct the image drifting, 100 nm gold nanoparticles (753688-25ML, Sigma-Aldrich) were used as fiducial markers in the imaging process. DNA imager stock solution was diluted and used in 1× PBS+500 mM NaCl at a working concentration of 2 nM.

### Image processing and quantitative comparison of ExRPath and DNA-PAINT in cultured neurons

2-dimensional DNA-PAINT images were rendered using Picasso in Python {Schnizbauer 2017} using a blur width of 0.0, oversampling of 9.477 (corresponding to a pixel resolution of 11.61 nm/pixel), and max density of 300. The oversampling parameter was calculated to match the pixel resolution given by the average measured expansion factor of ∼14x for ExRPath-processed cultured neurons in the 0.05x PBST condition. The expansion factor for ExRPath-process cultured neurons was calculated by measuring the distances between identical pairs of SMI structures in pre- and post-expansion fields of view, dividing the physical units in the post-expansion image by the physical units in the 150x confocal pre-expansion image, and averaging across the fields of view in the well (well 3 mean: 11.23x; well 3-2 calculated mean, 17.32x; overall mean: 14.274). The difference in measured expansion factor between the two fields of view in the same well is due to the use of a single 2-D slice for the pre-expansion 150x confocal image versus a maximum intensity projection of a stitched 3-D stack for the ExRPath image, which, depending on the orientation of the SMI structures, may distort relative distances between structures in different planes. Nevertheless, using the rounded well average expansion factor of 14x resulted in DNA-PAINT renderings that were properly scaled with ExRPath images (**Fig. 1b**). Post-expansion ExRPath stacks were first stitched either using Nikon’s built-in microscope stitching software or Fiji’s Image Stitching (Grid/Collection stitching {Preibisch 2009}), background subtracted using Fiji’s Rolling Ball algorithm with radius 50 pixels and collapsed to two dimensions using a maximum intensity projection. DNA-PAINT and ExRPath images were registered using a rigid transform implemented in custom MATLAB scripts. Images were first min-max normalized, passed through a 2-D Gaussian filter (sigma = 5), and binarized using Otsu’s thresholding method (MATLAB’s *graythresh*). The geometric transform was estimated using MATLAB’s configurations for multimodal intensity-based registration, with the following optimizer parameters: initial radius, 0.004, epsilon = 1.5x10^-4^, growth factor = 1.01, maximum iterations = 300. An initial rigid body transformation was estimated prior to calculation of the final transform using the same parameters. To eliminate small, high spatial frequency noise arising from gold nanoparticles for display (**Fig. 1b**), the registered ExRPath image was passed through a 2D median filter of radius 10 pixels and is displayed at 35% pixel saturation.

Root mean squared error (RMS) between ExRPath and DNA-PAINT images (**Fig. 1c**) was calculated as previously described {Chen 2015}. Briefly, a custom MATLAB script was used to implement a B-spline based non-rigid registration between pre and post expansion images, yielding vector fields for deformation within the images. These vector fields were then used to calculate root-mean-square length distortions across varying lengths. For plotting, RMS error values were binned into measurement length increments of 1um, and the mean RMS error within each bin is shown.

To assess the accuracy/resolution and distortion of ExRPath compared to DNA-PAINT, we calculated linear distortion of neurofilament staining in identical cultured-neuron samples imaged using the two technologies as previously described {Sarkar et al., Nat Biomedical Engineering 2022}, with a few modifications to account for the more diffuse nature of SMI staining compared to synaptic puncta staining as used in the original ExR validation. Briefly, small (400-1500 nm in linear extent) ROIs containing segments of neurofilament (SMI immunoreactivity) were manually segmented from DNA-PAINT and corresponding ExRPath images taken from the same fields of view in the same specimens. Following min-max normalization, gaussian filtration with sigma = 3, and intensity-based registration as previously described {Sarkar et al., Nat Biomedical Engineering 2022}, we calculated pixel-wise autocorrelation and cross-correlation as a function of shift distance. ROI pairs with less than 500 voxels of volume overlapping or nonzero volume in either ExRPath or DNA-PAINT volumes were excluded from analysis (listed sample size is after this exclusion). We then calculated the pairwise linear correlation coefficient between pixel intensity values in DNA-PAINT and ExRPath staining channels, or PAINT-PAINT or ExRPath-ExRPath staining channels for autocorrelation.

To calculate the half-maximal shift distance, we fit a third-degree polynomial (MATLAB’s ‘fit’ with ‘poly3’) to the correlation or autocorrelation function, and used the best-fit curve to estimate the shift distance at which the correlation or autocorrelation reached 50% of its maximum value.

Each of these calculations was repeated for each shift in the x- and y-dimensions. The mean differences in half-maximal shifts over x- and y-dimensions are plotted in Extended Data Fig. 1A,B.

## Single-shot 20ExM of human tissue slices (15ExMPath)

AcX treatment was performed as described above for the ExRPath protocol. Single-shot 20-fold Expansion Microscopy (20ExM) of human brain slices was performed following previously established methods {Wang, 2024, 20ExM https://www.nature.com/articles/s41592-024-02454-9#Sec8), with modifications optimized for human tissue specimens. In contrast to the original 20ExM protocol, which uses AX for protein anchoring, 15ExMPath employs AcX treatment.

To generate hydrophobic glass surfaces, coverslips were immersed in 0.2% (v/v) trichloro(octadecyl)silane (Fisher Scientific, AC147400250) in hexane for 90 seconds. Coverslips were then rinsed sequentially with 70% isopropanol and double-distilled water, dried at 37 °C, and gently wiped with a dry Kimwipe to remove residual silane.

The gelation solution was prepared by dissolving 0.522 g of sodium acrylate (SA) (AK Scientific, R624) in 1 ml of acidified Tris buffer, consisting of 10% (v/v) 1 M Tris-HCl (pH 8) and 20% (v/v) 1.2 M HCl in double-distilled water. Next, 7.5 µl of 10% (v/v) tetramethylethylenediamine (TEMED) (Sigma, T7024) and 900 µl of N,N-dimethylacrylamide (DMAA) (Sigma, 274135) were added. The mixture was vortexed until it became clear.

The gelation solution was placed on ice and degassed for 50 seconds using a gas dispersion tube (ChemGlass, CG-203-04) connected to a compressed nitrogen tank. After degassing, the solution was returned to room temperature (∼24 °C) and used for all subsequent gelation steps. To maintain an oxygen-free environment, all gelation components and tools—including the gelation solution, initiator solution (45 mg/ml potassium persulfate in ddH₂O), brain tissue slices, pipettes (P1000, P200, P20), pipette tips, humidified chamber, hydrophobic glass slides and coverslips, tweezers, transfer pipette, and 1.5-ml centrifuge tubes—were transferred into a glove bag (GlasCol, 108D X-17-17HG) filled with nitrogen. The glove bag was purged three to five times to remove residual oxygen.

Human brain slices were gently dried on hydrophilic coverglasses (Avantor, 48382-139) to ensure firm contact with the glass surface. Using hydrophobic coverglasses on both the bottom and top surfaces, as described in the original method, resulted in tissue tilting and poor imaging procefure; therefore, hydrophilic coverslips were used for the bottom surface. Inside the glove bag, 4 µl of initiator solution was added to 411 µl of gelation solution in a 1.5-ml centrifuge tube and mixed by inverting the tube five times. 50 µl of the mixture was applied to the tissue and incubated for 15 min in a humidified chamber. A hydrophobic coverslip was then placed on top to complete the gelation chamber. The sealed chamber was placed into an airtight humidified container, removed from the glove bag, and incubated at room temperature (∼24 °C) overnight (16–20 hours).

Following gelation, the portion of the gel containing tissue was excised and incubated in denaturation buffer (1 ml: 5% (v/v) SDS, 200 mM NaCl, 50 mM Tris pH 8.0, and 10 mg/ml DTT) overnight at 70 °C and 60 min at 121 °C and 30.5 psi (VWR, CLS-1191), instead of 1 h at 95 °C denaturation step used in the original 20ExM protocol for mouse brain tissue. Denatured gels were washed in 1× PBS five times for 15 min each. Note: termination of gelation and initiation of denaturation must be performed immediately after opening the airtight container; otherwise, the resulting gel may become fragile and exhibit a reduced expansion factor.

Immunostaining and imaging were performed according to the ExRPath protocol. However, due to the delicate nature of fully expanded 15ExMPath gels, final expansion was carried out directly within a glass-bottom well plate to avoid damage during transfer.

## Root mean square (RMS) error analysis of 15ExMPath

Root mean square (RMS) error analysis was assessed by confocal imaging of intact human tissue sections acquired both before and after expansion. Control human tissues (n=2) were immunostained with SMI (primary antibody, chicken host) followed by Alexa Fluor 546–conjugated anti-chicken secondary antibodies. Following the 15ExMPath protocol, the expanded samples were re-stained using the same antibody set. For analysis in the xy plane, post-expansion confocal images were Gaussian filtered, background-subtracted using Fiji’s rolling ball algorithm (radius = 50 pixels), and collapsed into two-dimensional images via maximum intensity projection. These processed post-expansion images were then rigidly registered to the corresponding pre-expansion images using Fiji (TurboReg, Scaled Rotation/Accurate/Manual). Subsequent nonrigid registration and calculation of deformation vector fields were performed in MATLAB. For xz and yz plane analyses, confocal z-stacks of the same brain regions were acquired and re-sliced into orthogonal views using Fiji’s orthogonal view tool, followed by Gaussian filtering. Both pre- and post-expansion images were subjected to rigid body registration in Fiji and nonrigid registration in MATLAB using the same workflow as applied for the xy plane analysis.

## Histology on Human Brain Tissue Samples

Human brain tissue histology and staining was performed as previously described in Mathys et al., Nature 2019 {Mathys 2019}. Fixed slices were mounted on slides. Antigen retrieval was performed by boiling the sections in citrate buffer for twenty minutes. Free floating sections were cooled and blocked with a buffer containing 0.3% Triton-X and 10% bovine serum albumin for 2 hours at room temperature, incubated with primary antibody overnight at 4°C, washed with

PBS-T, and incubated with secondary antibodies for 2 hours at room temperature. Lipofuscin autofluorescence was quenched using TrueBlack (Biotium 23007) according to the manufacturer’s protocol. Sections were subsequently washed with PBS mounted on slides. Slides were imaged using a Zeiss LSM 880 microscope with the same parameters for each image, and analyzed using Imaris software.

## 5XFAD Mouse Intracranial SARS-CoV-2 Spike Protein Injections

Mice were anesthetized with gaseous isoflurane immediately before surgery and kept on a heating pad throughout the procedure. A stereotaxic apparatus was used to inject at coordinates -1.8 mm anteroposterior, 2.2 mm lateral, relative to bregma. Holes were drilled with a dental drill bit. A microneedle loaded with SARS-CoV-2 S1 spike protein (Abcam) or vehicle control was inserted at these coordinates with 1.8 mm vertical. After waiting 5 minutes, 2.3 μL injections were administered at 100 nL/mm, and after waiting another 5 minutes, the microneedle was retracted and the incision was closed with surgical sutures. The animals recovered within 10 minutes and were singly-housed thereafter.

## Histology on Mouse Brain Tissue Samples

Samples analyzed via ExRPath were processed as described above. For conventional immunohistochemistry, the sections were incubated overnight at 4 °C in primary antibody in PBS with 0.3% Triton X-100 and 5% normal donkey serum. The sections were then washed with PBS room temperature (4 × 15 min), and incubated with secondary antibodies (dilution 1:2,000) and Hoechst (dilution 1:10000) for 12 hours at 4 °C. Slices were then washed with PBS at room temperature (4 × 15 min), before mounted on Fisherbrand Superfrost Plus microscope slides in ProLong Gold Antifade Mountant. Images were obtained using a confocal microscope (LSM 880; Zeiss) with a 20× objective.

## Imaris Quantifications

Imarisx64 8.1.2 (Bitplane, Zurich, Switzerland) was used to analyze IBA1-positive microglia clustering patterns, MHC-II signals and 6e10 intensity in 5XFAD mice with and without spike protein injections. To characterize microglia cell clustering, background was subtracted in Imaris and surface functions were applied to IBA1-positive signals. Clusters of cells within 15 μm regions of interest were quantified. MHC-II-positive signal and 6e10-positive signal were similarly processed in Imaris by applying surface functions, removing background, and counting the number of surfaces identified by the software. To characterize the extent of overlap between MHC-II and 6e10, surface functions were applied to the signals of overlap between MHC-II and 6e10 and the number of surfaces were quantified. Brain tissue sections from 3-5 mice for used for each quantification and each data point represents the average for one animal. Unpaired t-tests were applied to the quantification.

## Human iPSC-Derived Microglia Cultures

All human iPSCs were maintained at 37°C and 5% CO2, in feeder-free conditions in mTeSR1 medium (Cat #85850; STEMCELL Technologies) on Matrigel-coated plates (Cat # 354277; Corning; hESC-Qualified Matrix). iPSCs were passaged at 60–80% confluence using ReLeSR (Cat# 05872; STEMCELL Technologies) and reseeded 1:6 onto Matrigel-coated plates. The iPSC lines were generated by the Picower Institute for Learning and Memory iPSC Facility as first described {Lin 2018}. Microglia-like cells were differentiated using a previously established protocol {McQuade 2018}. In brief, 3e6 cells were then plated onto AggreWell 800 microwells (Cat# 34815; STEMCELL Technologies) for embryoid body (EB) induction. After 48 hours, EBs were seeded onto Matrigel-coated 6-well tissue culture plates at a density of 15-30 EBs per well. EBs were differentiated into hematopoietic progenitor cells (HPCs) using the STEMdiff Hematopoietic Kit (Cat#05310; STEMCELL Technologies). Non-adherent HPCs were collected, centrifuged at 300 x g, and resuspended in 1 mL of microglia differentiation media (MDM) containing a mixed composition of half DMEM/F12 (Cat#11330-057; Thermo Fisher Scientific) and half Neurobasal media (Cat# 21103049; Gibco) supplemented with IL-34 (Cat#200-34; PeproTech) and m-CSF (Cat#300-25; PeproTech) {McQuade et al., 2018}. Cells were plated in 6-well tissue culture plates at 200,000 cells per well and maintained in MDM for at least two weeks before experiments.

## iPSC-derived microglia infection and treatment

Cells were plated onto a Millipore eight-chamber glass slide, at a density of 250,000 cells per well. Cells were treated with either SARS-CoV-2 nucleocapsid, S1 spike protein, or S1 RBD domain (200 nM) for 48 hours. Aβ-42 peptide treatment was administered at 20 nM. Cells were fixed in 4% paraformaldehyde for 15 minutes prior to staining or harvested directly for bulk RNA-sequencing. SARS-CoV-2 USA-WA1/2020 (Gen Bank: MN985325.1) was generously provided by the Gehrke Laboratory, originally obtained from BEI Resources and expanded and tittered on Vero cells. Cells were infected in DMEM +2% FBS for 48 hrs using multiplicity (MOI) of 0.5 for infection. All sample infections, sample processing, and harvesting with infectious virus was performed in the BSL3 facility at the Ragon Institute.

## iPSC-derived microglia immunohistochemistry

Cells were fixed using 4% paraformaldehyde/PBS for 15 min at room temperature, washed three times with washing buffer (0.1% Tween-20/PBS, 5 min), and permeabilized with 1% Triton X-100/PBS for 30 min. Samples were incubated in blocking buffer (3% BSA, 2% goat serum in PBS) for 30 min, then incubated with various primary antibodies including guinea pig anti-IBA1 (1:500, Synaptic Systems), rabbit anti-P2RY12 (1:100, Millipore Sigma), and mouse anti-CD74 (1:500, Abcam), overnight at 4 °C on a shaker. After washing, samples were incubated with corresponding Alexa secondary antibodies (ThermoFisher) at 1:1000 for 1 hour at room temperature on a shaker. After washing, images were obtained using a confocal microscope (LSM 710; Zeiss).

## Antibodies

### Primary antibodies for immunostaining of tissue slices and ExRPath samples

| Target | Host | Vendor Information | Dilution |
| --- | --- | --- | --- |
| Neurofilament (SMI) | chicken | Abcam ab4680 | 1:200 |
| IBA1 | guinea pig | Synaptic Systems 234 004 | 1:500 |
| Abeta (6E10) | mouse | Biolegend 803001 | 1:500 |
| Abeta (D54D2) | rabbit | Cell Signaling Technology CST8243 | 1:200 |
| Abeta (12F4) | mouse | Biolegend SIG39142 | 1:200 |
| MHC Class II | mouse | EMD Millipore MABF33 | 1:250 |
| Spike | mouse | GeneTex GTX632604 | 1:200 |
| Spike | rabbit | Abcam ab272504 | 1:200 |
| CD13 | rabbit | Abcam ab108310 | 1:200 |
| PECAM-1 | sheep | Biotechne AF806 | 1:200 |
| GFAP | chicken | Abcam ab4674 | 1:200 |
| Bassoon | guinea pig | Synaptic Systems 141 004 | 1:200 |
| Myelin sheath (PLP) | rabbit | Abcam ab28486 | 1:200 |

## Bulk RNA-sequencing of iPSC-derived microglia

700 μL of EtOH and Trizol were added at a 1:1 ratio to each well, and cells were collected into tubes. The procedure was then followed according to Zymo Direct-Zol Microprep kit instructions. Approximately 500 ng of each sample was submitted for library prep (Kappa HiFi) and bulk sequencing performed by the BioMicro Center at MIT’s Department of Biology, using the NextSeq Illumina platform. Raw FASTQ data were mapped to a reference transcriptome derived from the GRCh38 human genome assembly and quantified using htseq {Putri et al., 2022}.

Differential gene expression testing was performed with DESeq2 {Love et al., 2014} with median-ratio count normalization, parametric dispersion estimation, and additional count normalization by variance-stabilizing transformation. Pathways were analyzed using the Gene Ontology database of molecular function processes (2019) from the Mayaan laboratory; https://maayanlab.cloud/Enrichr/#libraries).

## Isolation of nuclei from frozen post-mortem brain tissue

Nuclei were isolated from flash-frozen post-mortem brain tissue following a previously published protocol {Mathys 2019}. In brief, post-mortem brain tissue was gently homogenized in 700 µl homogenization buffer (320 mM sucrose, 5 mM CaCl_2_, 3 mM Mg(CH_3_COO)_2_, 10 mM Tris HCl pH 7.8, 0.1 mM EDTA pH 8.0, 0.1% IGEPAL CA-630, 1 mM β-mercaptoethanol, and 0.4 U/µl recombinant RNase inhibitor (Clontech)) in a BSL3 facility using the gentleMACS Tissue Dissociator. Homogenized tissue was filtered through a 40 µm cell strainer, mixed with an equal volume of working solution (50% OptiPrep density gradient medium (Sigma-Aldrich), 5 mM CaCl2, 3 mM Mg(CH3COO)2, 10 mM Tris HCl pH 7.8, 0.1 mM EDTA pH 8.0, and 1 mM β-mercaptoethanol) and loaded on top of an OptiPrep density gradient (750 µl 30% OptiPrep solution (30% OptiPrep density gradient medium,134 mM sucrose, 5 mM CaCl2, 3 mM Mg(CH3COO)2, 10 mM Tris HCl pH 7.8, 0.1 mM EDTA pH 8.0, 1 mM β-mercaptoethanol, 0.04% IGEPAL CA-630, and 0.17 U/µl recombinant RNase inhibitor) on top of 300 µl 40% OptiPrep solution (40% OptiPrep density gradient medium, 96 mM sucrose, 5 mM CaCl2, 3 mM Mg(CH3COO)2, 10 mM Tris HCl pH 7.8, 0.1 mM EDTA pH 8.0, 1 mM β-mercaptoethanol, 0.03% IGEPAL CA-630, and 0.12 U/µl recombinant RNase inhibitor). The nuclei were separated by centrifugation (5 min, 10,000 g, 4 °C). A total of 100 µl of nuclei was collected from the 30%/40% interphase and washed with 1 ml of PBS containing 0.04% BSA. The nuclei were centrifuged at 300g for 3 min (4 °C) and washed with 1 ml of PBS containing 0.04% BSA. Then the nuclei were centrifuged at 300g for 3 min (4 °C) and re-suspended in 100 µl PBS containing 0.04% BSA. The nuclei were counted and diluted to a concentration of 1,000 nuclei per microliter in PBS containing 0.04% BSA. All procedures were carried out on ice or at 4 °C.

## Droplet-based snRNA-seq

Libraries were prepared using the Chromium Single Cell 3′ Reagent Kits v3 according to the manufacturer’s protocol (10x Genomics). The generated snRNA-seq libraries were sequenced using NextSeq 500/550 High Output v2 kits (150 cycles) or NovaSeq 6000 S2 Reagent Kits.

## snRNA-seq data preprocessing

Gene counts were obtained by aligning reads to the GRCh38 genome using Cell Ranger software (v.3.0.2) (10x Genomics). To account for unspliced nuclear transcripts, reads mapping to pre-mRNA were counted. After quantification of pre-mRNA using the Cell Ranger count pipeline, the Cell Ranger aggr pipeline was used to aggregate all libraries (without equalizing the read depth between groups) to generate a gene-count matrix. The Cell Ranger 3.0 default parameters were used to call cell barcodes. We used SCANPY {Wolf 2018} to process and cluster the expression profiles and infer cell identities. We kept only protein coding genes and filtered out cells with over 20% mitochondrial or 5% ribosomal RNA, genes expressed in fewer than 3 cells, and cells with fewer than 100 genes expressed. We normalized COVID+ and control datasets separately, log1p transformed data, performed batch correction using ComBat {Johnson 2018}, and filtered the dataset to the top 5000 most variable genes. We then performed PCA (k=50), used Harmony followed by BB-kNN {Polanski 2020} across individuals for batch integration, and used the resulting cell-cell graph to calculate the low dimensional embedding of the cells (UMAP). We performed Leiden clustering on the final dataset of 53k cells to annotate them into seven major cell types using established PFC marker genes {Mathys 2019}.

## Analysis of snRNA-seq cell type composition

We calculated compositional differences in COVID+ vs. control individuals by modeling the number of cells of a certain cell type or subtype from a specific individual relative to the total number of cells using a quasi-binomial regression model. We used the emmeans package in R to assess significance of the regression contrasts and used p.adjust with the fdr method to adjust p-values.

## Differentially expressed genes in snRNA-seq

We performed differential expression (DE) analyses with the MAST and Nebula {He 2021} methods. We subset the tested genes to only genes present in over 20% of cells. We calculated and included in regressions the top 5 components of unwanted variation as calculated using RUV on the pseudo-bulk level data (at the level of individuals) and as well as covariates for each cell’s counts per gene and number of captured genes. For Nebula we used a poisson mixed-model on the count data with an offset of the log10 total counts per cell. For MAST, we normalized each cell to a total library size of 10,000 counts. We adjusted p-values for multiple testing in all cases by using the p.adjust function in R with the fdr method. For our final set of differential genes in each analysis, we took all genes that were significant (adjusted p-value > 0.05) and concordant in both the MAST and Nebula results. We separated genes DE in at least 3 cell types from cell type-specific DE genes.

## DEG and module pathway enrichments

We performed DEG enrichments for each differential expression run using the gprofiler2 package in R as unordered queries, with a p-value cutoff of 0.05, and using GO:BP, REAC, WP, KEGG, and CORUM as annotation sources, and kept enriched terms with fewer than 1000 genes.

## Statistics and reproducibility

The investigators were blinded when doing the experiments and running data analyses. No outliers were removed from analyses unless explicitly stated. All data were expressed as mean ± s.e.m. unless explicitly stated. We used GraphPad Prism (version 5.01, GraphPad Software) for data display and statistical analysis. We used unpaired two-tailed Student’s t-test and unpaired two-tailed Mann–Whitney rank-sum or Wilcoxon signed-rank test to compare two normally and non-normally distributed datasets, respectively, unless explicitly stated.

## Reporting summary

Further information on research design is available.

## Data Availability

The raw and analyzed datasets are available from the corresponding authors on reasonable request.

Code Availability

All accompanying scripts will be made publicly available on GitHub and Zenodo at the time of acceptance.

---

**Extended Data Figure 1:**
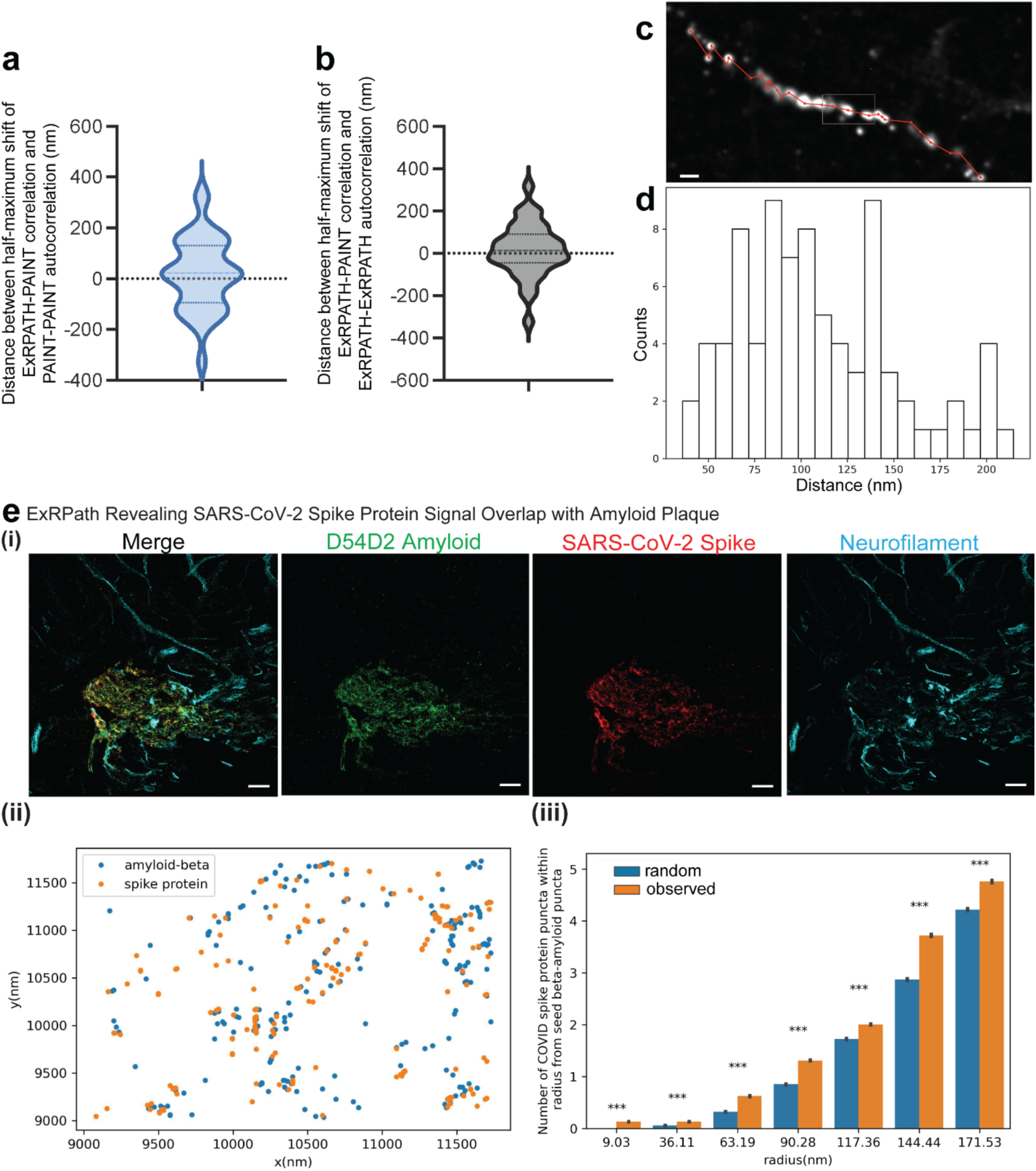
Quantification of autocorrelation between ExRPath and DNA-PAINT; quantification of a myloid-beta periodicity. a) Estimated population distribution (violin plot of density, with a dashed line at the median and dotted lines at the quartiles) of the shift (in nm) at which the correlation is half-maximal for PAINT-PAINT autocorrelation and ExRPath-PAINT correlation (calculated pixel-wise between intensity values normalized to the minimum and maximum of the image, see Methods; see **Extended Data Table 3** for statistics. n = 36 neurofilament ROIs from 2 fields of view from 1 well of cultured neurons from 1 culture batch). b), Same as (a) for ExRPath-ExRPath autocorrelation vs. ExRPath-PAINT correlation. c) Confocal image (maximum intensity projections) showing Aβ (white) clusters in COVID-decedent tissue from the dataset of Fig. 1g. Red lines connecting adjacent Aβ nanoclusters represent the distance between adjacent clusters (Scale bar, 100 nm). d) Histograms showing distances between adjacent Aβ (red) clusters along imaged segments of axons (n = 82 amyloid nanoclusters from 7 axonal segments from 2 human patients, mean = 109 nm). e) (i) ExRPath confocal images of max intensity projection showing co-localization of SARS-CoV-2 and Aβ (green: D54D2 Aβ, red: SARS-CoV-2, cyan: SMI neurofilament; scale bars, 2 µm); shown are Images from representative fields of view from two independent replicates (n = 3 tissue sections across n = 3 patients), (ii) Z projection of Aβ puncta (blue) and spike protein (orange) puncta represented by their local maximum in an ROI, (iii) the number of COVID spike protein puncta that are near to seed beta-amyloid puncta (represented in orange) is significantly greater than the number of randomly distributed spike proteins of the same total count (represented in blue) (one-tailed t-test, ***p-value < 0.001; see Extended Data Table 3 for statistics). The bar plot represents the mean values for each group while the gray error bars indicate the standard errors associated with the mean values.

**Extended Data Figure 2.**
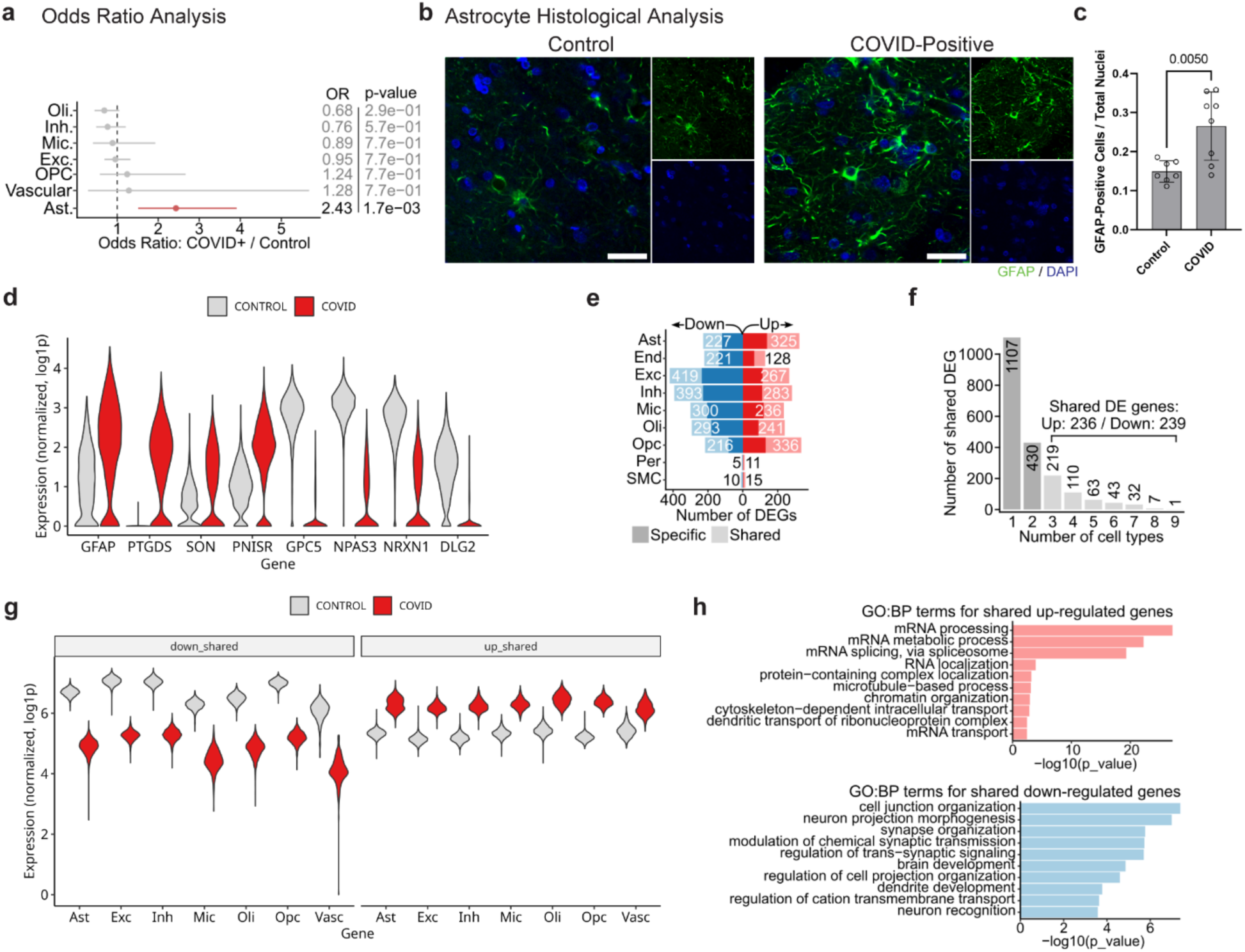
: Profiling inflammatory phenotypes in CNS of COVID+ decedents. **a)** Odds ratio of number of cells between COVID-positive and non-COVID individuals, **b)** immunohistochemistry for astrocyte marker GFAP in histological sections from non-COVID control (left) and COVID-positive (right) decedents (green: GFAP, blue: DAPI; scale bars, 50 µm), **c)** quantification of GFAP-positive cells between non-COVID control and COVID-positive groups (tissue from n = 8 individuals analyzed per group with n = 3 histological sections per individual; statistical analysis via t test; one outlier removed from control group that was more than 2.5 standard deviations outside of the mean), **d)** violin plots of top upregulated (left) and downregulated (right) DEGs in astrocytes between non-COVID control and COVID-positive decedent tissue at the cell level, **e)** number of differentially expressed genes (DEGs) altered in the COVID-positive condition for each individual cell type, **f)** number of shared DEGs segmented by the number of cell types displaying the shared DEG, **g)** violin plots displaying expression of shared DEGs across astrocytes (Ast), excitatory neurons (Exc), inhibitory neurons (Inh), microglia (Mic), oligodendrocytes (Oli), oligodendrocyte precursor cells (Opc), and vascular cells (Vasc) for shared downregulated DEGs (left) and upregulated DEGs (right) between non-COVID control and COVID-positive decedent tissue, and **h)** GO:BP terms for shared DEGs (3 or more cell types) for (top) upregulated or (bottom) downregulated genes.

**Extended Data Figure 3.**
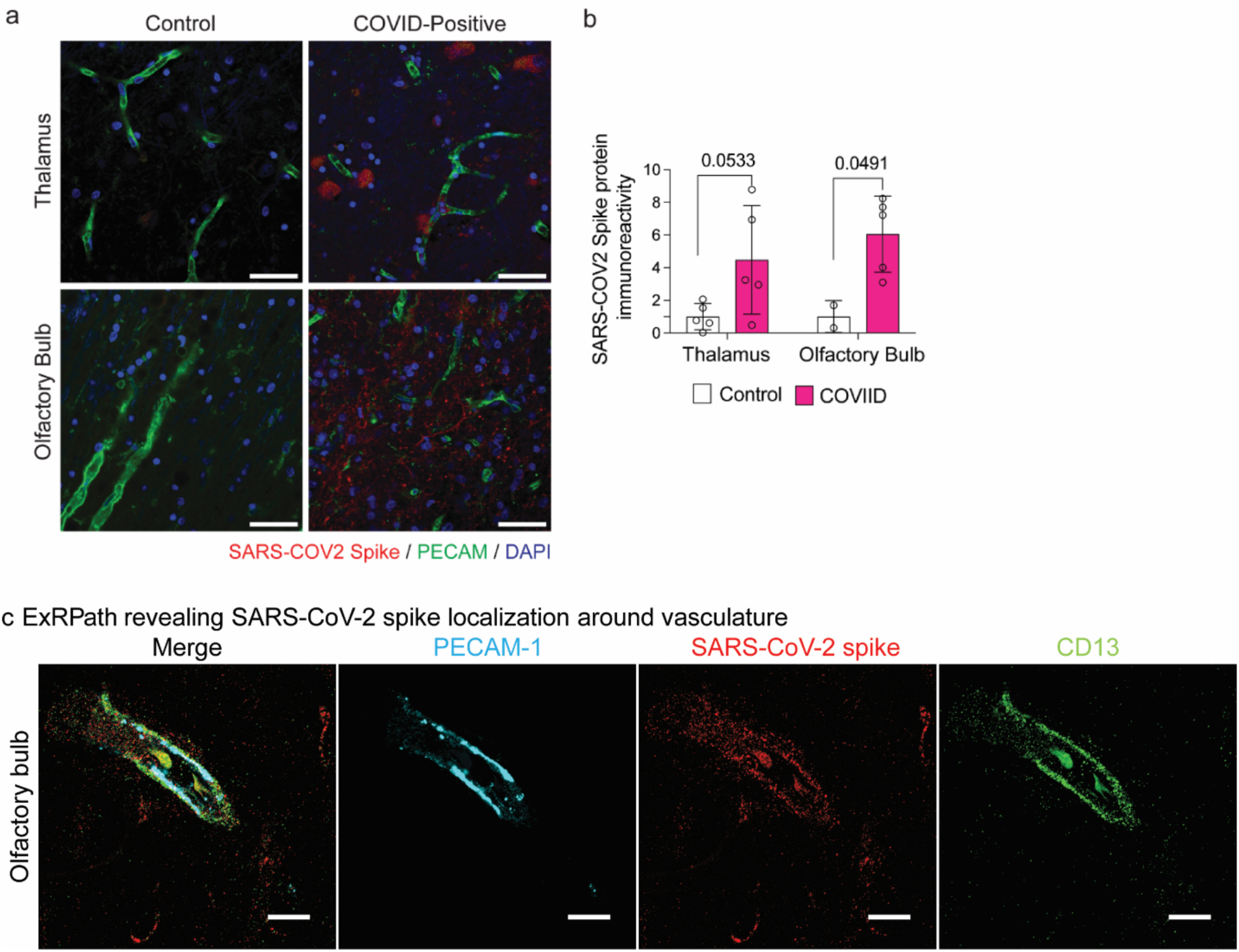
: SARS-CoV-2 localization in COVID-decedent brain tissue. **a)** SARS-CoV-2 immunoreactivity in thalamus (top) and olfactory bulb (bottom) tissue in non-COVID control (left) and COVID-positive decedent (right) tissue via conventional immunohistochemistry (red: SARS-CoV-2 spike protein, green: PECAM, blue: DAPI; scale bars, 50 µm) and **b)** quantification of SARS-CoV-2 immunoreactivity in tissues (n = 5 individuals except for olfactory bulb control n = 2 images for each of n = 4 tissue sections; statistical analysis via t test), **c)** ExRPath confocal images of max intensity projection revealing SARS-CoV-2 localization in the vasculature, characterized by double layered nanostructure between PECAM-1 (platelet endothelial cell adhesion molecule-1) and CD13 (brain pericytes marker) (cyan: PECAM-1, red: SARS-CoV-2 spike protein, green: CD13; scale bars, 2 µm). Shown are images from representative fields of view from two independent replicates (n = 3 tissue sections across n = 3 patients).

**Extended Data Table 1.** Statistical analysis for DNA-PAINT vs. ExRPath comparisons (Extended Data Fig. 1). Descriptive statistics for the population distribution of the difference between shift at the half-maximal correlation between ExRPath-PAINT correlation and PAINT-PAINT autocorrelation or ExRPath-ExRPath autocorrelation (nm).

|  | ExRPath-PAINT vs. PAINT-PAINT | ExRPath-PAINT vs. ExRPath-ExRPath |
| --- | --- | --- |
| Number of values | 36 | 36 |
| Minimum | -326.4 | -323.8 |
| Maximum | 334.2 | 316.2 |
| Range | 660.6 | 640 |
| Mean | 19.14 | 15.63 |
| Std. Deviation | 142.1 | 125 |
| Std. Error of Mean | 23.68 | 20.83 |
| 95% Confidence Interval of mean | [-28.93,67.21] | [-26.66,57.91] |

**Extended Data Table 2.** Patient Metadata. Value Coding 1 NCI: No cognitive impairment (No impaired domains) 2 MCI: Mild cognitive impairment (One impaired domain) and NO other cause of CI 3 MCI: Mild cognitive impairment (One impaired domain) AND another cause of CI 4 AD: Alzheimer’s dementia and NO other cause of CI (NINCDS PROB AD) 5 AD: Alzheimer’s dementia AND another cause of CI (NINCDS POSS AD) 6 Other dementia: Other primary cause of dementia

| GROUP | AGE | MALE | COGNITIVE STATUS | APOE GENOTYPE |
| --- | --- | --- | --- | --- |
| COVID | 96.09 | Female | 4 | 33 |
| COVID | 95.16 | Female | 2 | 34 |
| COVID | 90.32 | Female | 1 | 23 |
| COVID | 83.55 | Female | 4 | 33 |
| COVID | 94.23 | Female | 3 | 33 |
| COVID | 85.46 | Female | 1 | 34 |
| COVID | 97.09 | Male | 2 | 34 |
| COVID | 94.05 | Male | 2 | 33 |
| CONTROL | 87.20 | Female | 1 | 23 |
| CONTROL | 76.39 | Male | 1 | 33 |
| CONTROL | 90.44 | Female | 1 | 33 |
| CONTROL | 95.13 | Female | 1 | 33 |
| CONTROL | 77.31 | Female | 1 | 23 |
| CONTROL | 92.44 | Female | 2 | 34 |
| CONTROL | 94.32 | Female | 2 | 33 |
| CONTROL | 87.87 | Male | 1 | 33 |

**Extended Data Table 3.** . Full statistical analysis from Extended Data Fig. 1.

| radius | actual |  |  | random |  |  |  |  |  |
| --- | --- | --- | --- | --- | --- | --- | --- | --- | --- |
|  | mean | standard deviation | standard errors | mean | standard deviation | standard errors | t test statistics | p values | significance |
| **9.03nm** | 0.13 | 0.34 | 0.003 | 0 | 0.04 | 0.000 | 39.500 | 0.0 | *** |
| **36.11nm** | 0.13 | 0.34 | 0.003 | 0.06 | 0.24 | 0.002 | 18.487 | 9.302e-67 | *** |
| **63.19nm** | 0.62 | 0.71 | 0.007 | 0.32 | 0.58 | 0.006 | 33.938 | 1.641e-258 | *** |
| **90.28nm** | 1.31 | 1.03 | 0.010 | 0.85 | 0.93 | 0.009 | 33.897 | 2.093e-241 | *** |
| **117.36nm** | 2.01 | 1.32 | 0.013 | 1.73 | 1.36 | 0.013 | 15.221 | 1.103e-258 | *** |
| **144.44nm** | 3.72 | 1.93 | 0.019 | 2.87 | 1.69 | 0.017 | 33.969 | 1.418e-258 | *** |
| **171.53nm** | 4.76 | 2.29 | 0.022 | 4.22 | 2.05 | 0.020 | 18.013 | 3.911e-68 | *** |

## Notes

### Competing Interest Statement

The authors have declared no competing interest.

